# Leveraging host-genetics and gut microbiota to determine immunocompetence in pigs

**DOI:** 10.1101/2021.06.08.447584

**Authors:** Yuliaxis Ramayo-Caldas, Laura M. Zingaretti, David Pérez-Pascual, Pamela A. Alexandre, Antonio Reverter, Toni Dalmau, Raquel Quintanilla, Maria Ballester

## Abstract

The aim of the present work was to identify microbial biomarkers linked to immunity traits and to characterize the contribution of host-genome and gut microbiota to the immunocompetence in healthy pigs. To achieve this goal, we undertook a combination of network, mixed model and microbial-wide association studies (MWAS) for 21 immunity traits and the relative abundance of gut bacterial communities in 389 pigs genotyped for 70K SNPs. The heritability (h^2^; proportion of phenotypic variance explained by the host genetics) and microbiability (m^2^; proportion of variance explained by the microbial composition) showed similar values for most of the analyzed immunity traits, except for both IgM and IgG in plasma that were dominated by the host genetics, and the haptoglobin in serum which was the trait with larger m^2^ (0.275) compared to h^2^ (0.138). Results from the MWAS suggested a polymicrobial nature of the immunocompetence in pigs and revealed associations between pigs gut microbiota composition and 15 of the analyzed traits. The lymphocytes phagocytic capacity (quantified as mean fluorescence) and the total number of monocytes in blood were the traits associated with the largest number of taxa (6 taxa). Among the associations identified by MWAS, 30% were confirmed by an information theory network approach. The strongest confirmed associations were between *Fibrobacter* and phagocytic capacity of lymphocytes (r=0.37), followed by correlations between *Streptococcus* and the percentage of phagocytic lymphocytes (r=-0.34) and between *Megasphaera* and serum concentration of haptoglobin (r=0.26). In the interaction network, *Streptococcus* and percentage of phagocytic lymphocytes were the keystone bacterial and immune-trait, respectively. Overall, our findings reveal an important connection between immunity traits and gut microbiota in pigs and highlight the need to consider both sources of information, host genome and microbial levels, to accurately characterize immunocompetence in pigs.

## Introduction

The pig industry has a considerable socio-economical value representing around 35% of the total meat produced worldwide [1] and being the most popular meat for consumption [2]. The intensification of pig production coupled with the ban on in-feed use of antibiotics has led to a deterioration of the health status of pig farms. In addition, the current emergence of antibiotic resistance and society demands for healthier products and environmentally responsible livestock systems, has motivated to explore relevant approaches for pig and other livestock breeding programs, to improve robustness and disease resistance [3].

The implementation of breeding programs to select animals according to their robustness presents several challenges and levels of complexity. One of the most relevant milestones is the identification of selection criteria that combine functional traits with those of immunocompetence. These complex traits are driven by several physiological and behavioral mechanisms that in turn are determined by genetic and environmental factors. Regarding the genetic determinism of immunocompetence, several studies in pigs acknowledged medium to high heritability estimates [4–9] and reported genomic regions and candidate genes associated with phenotypic variation of health-related traits [9–15].

Over the past few years, multiple studies highlighted the relevant role of the gut microbiota composition in the homeostasis and function of the mammalian immune system [16–19]. Gut microbiota can regulate host-immunity through both direct mechanisms like translocation of bacteria and their components (i.e. metabolites), or mediate indirect process such as T-cell polarization and the regulation of immune cell trafficking [18]. Commensal gut populations modulate hosts’ immune responses, which in turn can modify the microbiota composition to maintain gut homeostasis [20, 21]. Recently, polymorphisms located in immune genes associated with the abundance of microbial communities have been reported [22–25]. Furthermore, it has been suggested that the pattern recognition receptors, which are proteins capable of recognizing molecules frequently associated with pathogens, may have evolved to mediate the bidirectional crosstalk between microbial symbionts and their hosts [26]. This has resulted in a mutualistic and symbiotic partnership between the immune system and these commensal microorganisms [27]. Therefore, the immune system not only protects the host from pathogens but can also modulate, and is itself modulated, by beneficial microbes.

Considering the relevant interplay between gut microbiota and host immunity, a better understanding of the role of gut microbiota in the immunocompetence determination in pigs could greatly assist in the implementation of selection programs to improve robustness and disease resistance simultaneously. The present work aimed to identify microbial biomarkers linked to immunity traits and to estimate the contribution of host-genome and gut microbial communities to the immunocompetence in healthy pigs.

## Material and Methods

### Ethics Statement

All experimental procedures were performed according to the Spanish Policy for Animal Protection RD53/2013, which meets the European Union Directive 2010-63-EU about the protection of animals used in experimentation. The experimental protocol was approved by the Ethical Committee of the Institut de Recerca i Tecnologia Agroalimentàries (IRTA).

### Animal Samples

Samples employed in this study are a subset of pigs reported in Ballester et al. [9] and Reverter et al. [25]. A total of 405 weaned piglets (204 males and 201 females) from a commercial Duroc pig line were used. The pigs were distributed in six batches obtained from 132 sows and 22 boars. All animals were raised on the same farm and fed *ad libitum* a commercial cereal-based diet.

### Immunity and hematological traits

Details of the sampling and laboratory processing have been reported [9]. In brief, blood and saliva samples were collected from all 405 piglets at 60 ± 8 days of age. Blood samples in 4 ml EDTA tubes were used to measure the hemograms (Laboratory Echevarne, Spain; Barcelona). Saliva was collected with Salivette tubes (Sarstedt S.A.U., Germany) according to the protocols recommended by the manufacturer. Blood samples for serum were collected in 6 mL tubes with gel serum separator and centrifuged at 1600 g for 10 min at RT. Plasma was collected from the sampled blood in 6 ml heparinized tubes and centrifuged at 1300 g for 10 minutes at 4°C. Plasma and serum samples were collected, aliquoted, and stored a −80°C. The following haematological parameters were included in this study: total number of eosinophils (EO), leukocytes (LEU), lymphocytes (LYM) and neutrophils (NEU) in blood. Analyzed immunity parameters included immunoglobulins (IgA, IgG and IgM) concentrations in plasma; C-reactive protein (CRP), Haptoglobin (HP) and Nitric Oxide (NO) concentrations in serum; and IgA concentration in saliva (IgAsal). Gamma-delta T cells (γδ T cells) were separated from heparinised peripheral blood by density-gradient centrifugation with Histopaque-1077 (Sigma, Spain). Phagocytosis assay was carried out in heparinized whole blood samples incubated with fluorescein (FITC)-labelled opsonized *Escherichia coli* bacteria using the Phagotest kit (BD Pharmigen, Spain) as indicated in the manufacturer’s protocol. The following phagocytosis traits were used: percentage of total phagocytic cells (PHAGO_%); percentage of phagocytic cells among granulocytes (GRANU_PHAGO_%), monocytes (MON_PHAGO_%) and lymphocytes (LYM_PHAGO_%); mean fluorescence in FITC among the total phagocytic cells (PHAGO_FITC); and mean fluorescence in FITC among the granulocytes (GRANU_PHAGO_FITC), monocytes (MON_PHAGO_FITC) and lymphocytes (LYM_PHAGO_FITC) that phagocyte.

### DNA extraction, sequencing and bioinformatics analysis

Simultaneous with blood and saliva samples, fecal samples were collected from all 405 piglets. DNA was extracted with the DNeasy PowerSoil Kit (QIAGEN, Hilden, Germany) following manufacturer’s instructions. Extracted DNA was sent to the University of Illinois Keck Center for Fluidigm sample preparation and paired-end (2 × 250 nt) sequencing on an Illumina NovaSeq (Illumina, San Diego, CA, USA). The 16S rRNA gene fragment was amplified using the primers V3_F357_N: 5’-CCTACGGGNGGCWGCAG-3’ and V4_R805: 5’-GACTACHVGGGTATCTAATCC-3’. Sequences were analysed with *Qiime2* [28]; barcode sequences, primers and low-quality reads (Phred scores of <30) were removed. The quality control also trimmed sequences based on expected amplicon length and removed chimeras. Afterwards, sequences were processed into Amplicon Sequences Variants (ASVs) at 99% of identity. Samples with less than 10,000 reads were excluded and ASVs present in less than three samples and representing less than 0.001% of the total counts were discarded. ASVs were classified to the lowest possible taxonomic level based on a primer-specific trained version of GreenGenes Database [29].

### Genotype data

A total of 390 out of 405 animals were genotyped using the Porcine 70 k GGP Porcine HD Array (Illumina, San Diego, CA) containing 68,516 single nucleotide polymorphisms (SNPs). The quality control excluded SNPs with minor allele frequencies <5%, rates of missing genotypes above 10%, and SNPs that did not map to the porcine reference genome (Sscrofa11.1 assembly). Consequently, 42,641 SNPs were retained for subsequent analysis.

### Microbiability and heritability estimation

Heritability (h^2^), i.e. the proportion of variance explained by the host genetics, and microbiability (m^2^), i.e. the proportion of variance explained by the microbial composition, were estimated for each immunity trait based on a mixed-model as follows:

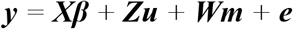

where ***y*** is the *n*-dimensional vector containing the individual phenotypes for the immune trait under consideration; ***β*** is the vector of fixed effects, containing the general intercept, the sex effect (two levels), and batch effect (six levels) for most traits but data of laboratory analysis (12 levels, two by batch) for phagocytosis-related traits; ***u*** is the vector containing the host genetic random effect from each individual; ***m*** is the vector of the animal’s microbiome random effect; ***X***, ***Z*** and ***W*** are, respectively, the incidence matrices correspondent to ***β***, ***u*** and ***m***; and **e** is the vector of residual terms.

Assuming independence between random effects, the following distributions were considered: *u* ~*N*(0,G, σ^2^_u_), where σ^2^_u_ is the host genetic effects variance and *G* is the genomic relationship matrix between individuals, computed following [30], i.e., 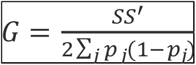 being ***S*** the matrix that contains the centered individual genotype for the 42,641 SNPs (columns) of each individual (rows), and *p_j_* is the frequency of the minimum allele of the *j^th^* SNP; *m* ~*N*(0,B, σ^2^_m_), where σ^2^_m_ is the microbial effects variance and ***B*** the microbial relationship matrix computed following [31], i.e., 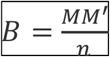, being ***M*** the matrix containing the scaled after a previous cumulative sum scaling normalization of the ASV abundances (columns) for each individual microbiome (rows) and *n* the total number of ASVs; and finally 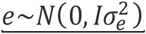, where 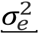 is the error variance.

The model parameters for each immunity trait were estimated by a Bayesian approach, using the Bayes Ridge Regression model from BGLR package [32]. We used a Gibbs sampler with 30,000 iterations and a burn-in of 3,000 rounds. The ‘heritability’ 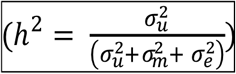 and ‘microbiability’ 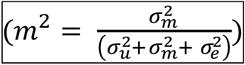 were estimated from the mean of the posterior distributions [33].

### Microbial Wide Association Study

We performed a Microbial Wide Association Study (MWAS) using a multi-ASV association method that combines all the ASVs in a single model:

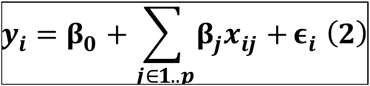

Given a trait ***y_i_*** measured in *n* individuals and a matrix ***X*** containing relative abundances of *p* taxa from a microbial community, here the ASVs effects were treated as draws from normal distributions as in any Bayesian Ridge Regression approach [32].

Following the approach of Legarra et al [34], Bayes Factor (BF) for the effect of each taxa can be derived as the ratio of probabilities 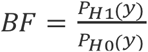, where H1 means “the *j*-genus has some effect” and H0 “the *j*-genus has no effect”. The calculations from the posterior distribution are very simple since both probabilities 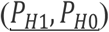 are normal density.

### Network between microbial and immunity traits

To better understand the relationship between microbial communities and immunity traits we implemented PCIT [35], a network-based approach that combines partial correlation coefficient with information theory to identify significant correlations between each possible combination of clr-transformed bacterial abundance and the immune-traits [35]. PCIT tests all possible 3-way combinations in the dataset and only keeps correlations between traits if they are significant and independent of the association of another features. To reduce the complexity of the resulting network, from the PCIT significant connections, we kept only the ones involving one immune-trait and one genus (i.e. genus-genus and trait-trait interactions were no represented).

## Results

In this study, 16S rRNA gene sequences, host genotype information and immune traits from 389 Duroc pigs were analyzed to estimate both host genomes and gut microbiota contribution to the porcine immunocompetence, and to identify microbial biomarkers linked to immunity traits. Table 1 summarizes the immunity traits and their descriptive statistics used in the present study. Regarding 16S rRNA gene sequences, after quality control, a total of 2,055 Amplicon Sequences Variants (ASVs) and 68 genera were detected. The dominant bacterial phyla were Bacteroidetes and Firmicutes, and the most abundant genera were *Prevotella*, *Lactobacillus*, *Treponema*, *Roseburia* and *Ruminococcus* (Supplementary figure 1)

**Table 1.** Descriptive statistics, mean, standard deviation (SD) and coefficient of variation (CV) of the 21 analysed traits.

| Trait | Abrev | Mean | SD | CV |
| --- | --- | --- | --- | --- |
| C-reactive protein in serum (µg/ml) | CRP | 176.69 | 128.61 | 0.73 |
| Eosinophils count n/µL | EO | 406.31 | 191.04 | 0.47 |
| γδ T-lymphocyte subpopulation (%) | γδ T-cells | 7.83 | 4.94 | 0.63 |
| Granulocytes phagocytosis FITC | GRANU_PHAGO_FITC | 5.15 | 0.42 | 0.08 |
| Granulocytes phagocytosis (%) | GRANU_PHAGO_% | 91.70 | 3.85 | 0.04 |
| Haptoglobin in serum (mg/ml) | HP | 1.03 | 0.67 | 0.65 |
| IgA in plasma (mg/ml) | IgA | 0.65 | 0.31 | 0.48 |
| IgA in saliva (mg/dl) | IgAsal | 5.13 | 3.12 | 0.61 |
| IgG in plasma (mg/ml) | IgG | 12.64 | 4.85 | 0.38 |
| IgM in plasma (mg/ml) | IgM | 2.27 | 0.79 | 0.35 |
| Leukocytes count n/µL | LEU | 20,350.15 | 6948.01 | 0.34 |
| Lymphocytes count n/µL | LYM | 12,422.40 | 4497.03 | 0.36 |
| Lymphocytes phagocytosis FITC | LYM_PHAGO_FITC | 3.16 | 0.13 | 0.04 |
| Lymphocytes phagocytosis (%) | LYM_PHAGO_% | 6.06 | 4.02 | 0.66 |
| Monocytes phagocytosis FITC | MON_PHAGO_FITC | 3.92 | 0.25 | 0.06 |
| Monocytes phagocytosis (%) | MON_PHAGO_% | 49.48 | 9.72 | 0.20 |
| Monocytes count n/μL | MONOCITOS_MM | 548.42 | 280.43 | 0.51 |
| Nitric oxide in serum (μM) | NO | 206.45 | 79.83 | 0.39 |
| Phagocytosis FITC | PHAGO_FITC | 4.71 | 0.33 | 0.07 |
| Phagocytosis (% cells) | PHAGO_% | 42.94 | 8.37 | 0.19 |
| Neutrophils count n/μL | NEU | 6985.62 | 3316.61 | 0.47 |

### Heritability and microbiability of immunity traits

Posterior estimates of *h^2^* and *m^2^* for the 21 health-related traits can be shown in Figure 1 and Supplementary Table 1. Posterior means of h^2^ in the analyses considering microbiota contribution reached low to medium values (from 0.138 to 0.359), but posterior probability of h^2^ being superior to 0.1 was in all cases above 0.82. Similarly, estimated m^2^ reached values between 0.152 and 0.276, and the probability of being above 0.1 was above 0.85 for all immunity and hematological traits (Supplementary Table 1). Among analysed traits, IgG and IgM in plasma showed the highest genetic determinism (h^2^= 0.316 and 0.359), whereas microbiota contribution was below 0.18. Conversely, the Hp concentration in serum showed the highest microbial effect (m^2^=0.276), accompanied by the lowest h^2^ estimate (h^2^= 0.138). Considering the joint effects of host-genome and gut microbiota, these two sources of variation explained from 29.9% to 51.7% phenotypic variance of the analysed immunity and hematological traits. To be noted, in the 76% (16/21) of these traits the *h^2^* and *m^2^* estimates reaching relatively similar values (Figure 1).

**Figure 1.**
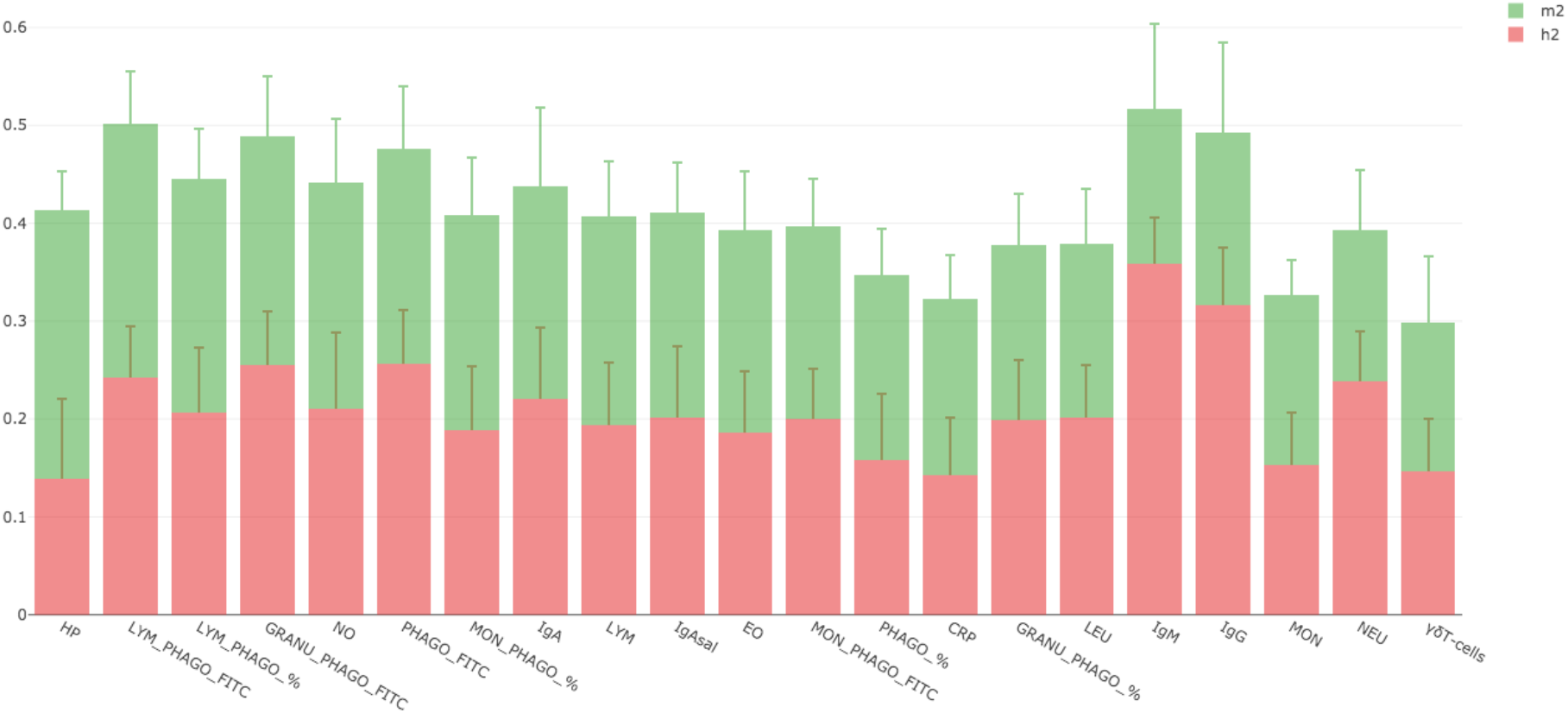
Percentage of phenotypic variance explained by the host-genetic (red points) and the gut microbial composition (green points) for most relevant immunity traits.

### Associations between microbial genera and immunity traits

Results from the MWAS reported some putative associations between bacterial genera abundance and health-related traits (Table 2). In particular, 15 out of the 21 immunity traits were associated with at least one microbial genus (Table 2). The strongest association was observed between the relative abundance of *Chlamydia* and the profile of LYM_PHAGO_FITC, followed by *Streptococcus* linked to LYM_PHAGO_% and *Peptococcus* associated with LYM_PHAGO_FITC. In addition, several genera showed multiple associations with numerous immunity traits: *Desulfovibrio*, *Oribacterium* and *Chlamydia* (4 traits) followed by *Oxalobacter* and *Parabacteroides* (3 traits), *Peptococcus* and *Streptococcus* (2 traits). As far as the analysed phenotypes, those traits showing the highest number of associations with different bacterial taxa were: LYM_PHAGO_FITC and MON (6 taxa); LYM_PHAGO_%, EOS, GRANU_PHAGO_FITC (4 taxa) and total number of LEU (3 taxa). Meanwhile, only four out of the 15 immunity traits analysed were linked with only one genus (Table 2).

**Table 2.** Results from the microbial-wide association studies

| Names | Description_trait | Genus | BPcum |
| --- | --- | --- | --- |
| MON | Monocytes count n/μL | *Anaerobiospirillum* | 2.676 |
| LYM_PHAGO_FITC | Lymphocytes phagocytosis FITC | *Bacteroides* | 2.536 |
| IgG | IgG in plasma (mg/ml) | *Bulleidia* | 2.313 |
| GRANU_PHAGO_FITC | Granulocytes phagocytosis FITC | *Campylobacter* | 6.556 |
| MON | Monocytes count n/μL | *Catenibacterium* | 2.952 |
| EO | Eosinophils count n/μL | *Chlamydia* | 5.292 |
| IgAsal | IgA in saliva (mg/dl) | *Chlamydia* | 2.364 |
| LYM_PHAGO_FITC | Lymphocytes phagocytosis FITC | *Chlamydia* | 11.246 |
| LYM_PHAGO_% | Lymphocytes phagocytosis (%) | *Chlamydia* | 3.088 |
| EO | Eosinophils count n/μL | *Desulfovibrio* | 3.369 |
| LEU | Leukocytes count n/μL | *Desulfovibrio* | 2.085 |
| LYM_PHAGO_FITC | Lymphocytes phagocytosis FITC | *Desulfovibrio* | 2.852 |
| MONOCITOS_MM | Monocytes count n/μL | *Desulfovibrio* | 2.427 |
| LYM_PHAGO_FITC | Lymphocytes phagocytosis FITC | *Fibrobacter* | 3.829 |
| HP | Haptoglobin in serum (mg/ml) | *Megasphaera* | 3.582 |
| MON_PHAGO_FITC | Monocytes phagocytosis FITC | *Mitsuokella* | 2.788 |
| CRP | C-Reactive Protein in serum (ug/ml) | *Mucispirillum* | 2.588 |
| LYM_PHAGO_% | Lymphocytes phagocytosis (%) | *Oribacterium* | 3.089 |
| MON_PHAGO_FITC | Monocytes phagocytosis FITC | *Oribacterium* | 2.673 |
| PHAGO_% | % of total phagocytic cells | *Oribacterium* | 2.009 |
| NEU | Neutrophils count n/μL | *Oribacterium* | 2.478 |
| LEU | Leukocytes count n/μL | *Oxalobacter* | 2.816 |
| MON | Monocytes count n/μL | *Oxalobacter* | 2.320 |
| NEU | Neutrophils count n/μL | *Oxalobacter* | 3.216 |
| IgG | IgG in plasma (mg/ml) | *Paludibacter* | 3.018 |
| HP | Haptoglobin in serum (mg/ml) | *Parabacteroides* | 2.018 |
| LYM_PHAGO_FITC | Lymphocytes phagocytosis FITC | *Parabacteroides* | 2.219 |
| MON_PHAGO_FITC | Monocytes phagocytosis FITC | *Parabacteroides* | 5.158 |
| LYM_PHAGO_FITC | Lymphocytes phagocytosis FITC | *Peptococcus* | 7.497 |
| PHAGO_% | % of total phagocytic cells | *Peptococcus* | 2.288 |
| GRANU_PHAGO_FITC | Granulocytes phagocytosis FITC | *rc4.4* | 2.156 |
| LEU | Leukocytes count n/μL | *rc4.4* | 2.619 |
| LYM | Lymphocytes count n/μL | *rc4.4* | 3.621 |
| MON | Monocytes count n/μL | *RFN20* | 3.022 |
| GRANU_PHAGO_FITC | Granulocytes phagocytosis FITC | *Sphaerochaeta* | 2.686 |
| γδ T-cells | γδ T-Lymphocyte subpopulation | *Streptococcus* | 2.444 |
| LYM_PHAGO_% | Lymphocytes phagocytosis (%) | *Streptococcus* | 8.484 |
| EO | Eosinophils count n/μL | *Succinivibrio* | 4.814 |
| MON | Monocytes count n/μL | *Succinivibrio* | 2.439 |
| LYM PHAGO % | Lymphocytes phagocytosis (%) | *Treponema* | 2.187 |

### Gut microbial and host-immune interaction network

The interplay between microbial and health-related traits was also inferred through a network comprised of 63 nodes (42 genera and 21 immunity traits) and 86 edges (significant connections) in which only the significant interactions between a bacterial genus and an immunity trait were considered (Figure 2). The topological evaluation of the network highlights LYM_PHAGO_% as the most connected trait, followed by IgAsal, NEU and Hp. Meanwhile, at microbial level, *Streptococcus* was the most connected genus followed by *Acidaminococcus*, *Desulfovibrio* and *Blautia*. The network approach confirmed 30% (12/40) of the associations identified by the MWAS (Figure 2). The strongest confirmed correlation was between *Fibrobacter* and LYM_PHAGO_FITC (r=0.37) followed by correlations between *Streptococcus* and LYM_PHAGO_% (r=-0.34) and between *Megasphaera* and Hp (r=0.26). To be noted, *Streptococcus* and LYM_PHAGO_% that showed the strongest confirmed association in the MWAS were highly in the interaction-network as the keystone bacterial and immunity trait, respectively.

**Figure 2.**
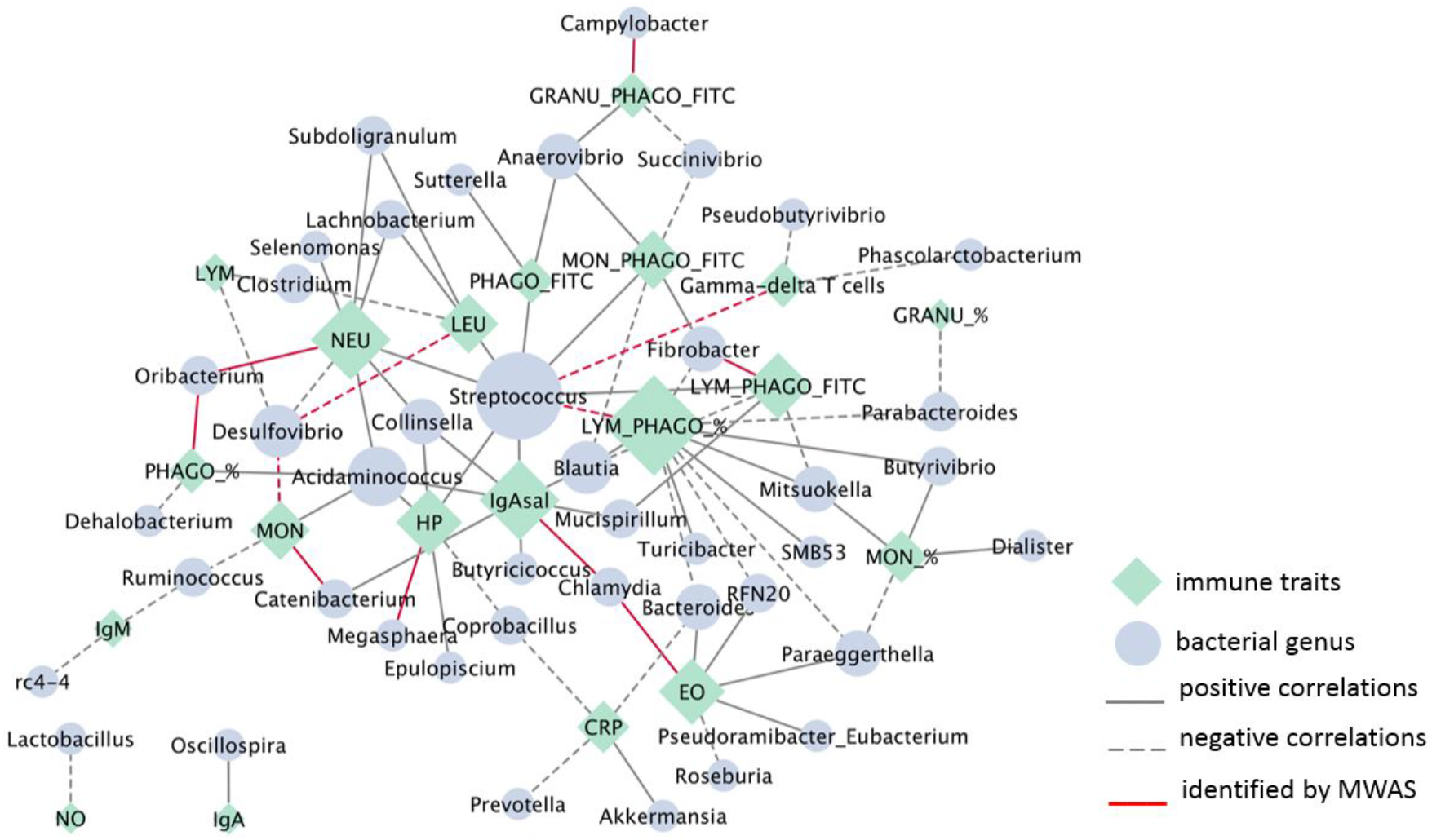
Microbial - health-related traits network. Green diamond nodes correspond to immunity traits (n=21) and blue ellipse nodes correspond to microbial genera (n=42). Node sizes are relative to their topological degree (number of connections) and edges are continuous or dashed to represent positive or negative correlations, respectively. Relationships previously identified by MWAS are highlighted in red.

## General Discussion

### Host-genome and gut microbiota contribution to porcine immunocompetence

We report the first study that aimed to dissect the joint contribution of the host genome and the gut microbiota to the immunocompetence in healthy pigs. Estimates of microbiability pointed out significant microbial effects on most immunity and hematological traits, ranging between 15% and 27% of total phenotypic variance. Effects of microbiota resulted particularly relevant for Hp concentrations in serum, followed by the parameters related to phagocytosis of lymphocytes. Regarding genomic heritabilities of these traits, they reached low to moderate values and were substantially lower compared to the medium to high h^2^ previously obtained in the same Duroc population [9] for all traits but MON and MON_PHAGO_%. A dramatic decrease of the estimated host genetic effects was observed for γδ T cells, but also for EOS and NEU counts and immunoglobulins concentrations in plasma, despite IgM and IgG variability seemed dominated by host genetics and showed the highest h^2^ among analysed traits. These results would call into question the high genetic determinism of the global immunocompetence in pigs reported in previous studies [6, 7, 9]. However, it should be considered that the limited sample size joint with the likely similarity between close relatives (particularly between full-sibs) in their microbiota profiles makes plausible that the model could not separate adequately genetic from microbiota effects.

### Microbial signatures associated with immunity traits

In the present study, we implemented a combination of MWAS and network approaches to pinpoint microbial signatures associated with immunity traits, revealing some interesting associations between the composition of the pig gut microbiota and the host immunity traits. Remarkably, lymphocyte phagocytosis traits were among the most connected and associated traits to the highest number of taxa and were also central nodes in the network. The strongest confirmed association involved *Fibrobacter* relative abundance in gut microbiota and the host phagocytosis capacity of lymphocytes, which were positively correlated (r=0.37). *Fibrobacter* genus is composed of strictly anaerobic bacteria with cellulolytic capacity capable of degrading complex plant fiber [36] and it has been associated with better feed efficiency in pigs [37, 38]. Conversely, the relative abundance of *Streptococcus* showed an opposite association with the percentage of phagocytic lymphocytes (r=-0.34). *Streptococcus* was also the keystone taxa in the network. In pigs, some *Streptococcus* species are important opportunistic pathogens such as *Streptococcus suis*, which abundance increased in the stomach and small intestine after weaning [39]. Piglets with high intestinal concentrations of *S. suis* can serve as a source of transmission and infection between animals and farms (reviewed in [39]). In general *Streptococcus* are less abundant in more-feed efficient pigs [37], although there are also evidences of the immunomodulatory properties of member of *Streptococcus* genus [40, 41].

Several studies in mammals have demonstrated that B cells have a significant phagocytic capacity, being able to phagocytose particles including bacteria [42–44]. Most important has been the demonstration of the efficient capability of these cells to present antigen from phagocytosed particles to CD4^+^ T cells [42–44], acting as a bridge that link innate with adaptive immunity. Therefore, considering the inferred high connection of these phagocytosis phenotypes with gut microbiota, we could hypothesize that, as other antigen-presenting cells such as dendritic cells or macrophages, the phagocytic lymphocytes seem to be relevant to maintain immune tolerance to the normal gut microbiota, being also relevant to control the abundance of opportunistic pathogens. B-cells also produce secretory IgA, the most abundant secreted isotype in mammals and a key element to maintain ‘homeostatic immunity’[45]. Secretory IgA was among the most connected traits in the network being positively associated with several taxa. Among them, the genus *Blautia* is of particular interest due to its potential role modulating inflammatory and metabolic diseases, with potential beneficial effects for the host [46]. Therefore, similar to phagocytic lymphocytes, the interplay between secretory IgA levels and the abundance of different taxa in our animals may regulate the ecological balance of commensal bacteria and the development of Ig-A secreting cells. Neutrophils were also positively correlated with gut microbiota profiles. A systemic immunomodulation of neutrophils by intestinal microbiota has been demonstrated [47], and a crosstalk between NEU and gut microbial composition has been also documented [48]. Our results confirmed a positive association of *Oribacterium* abundance with the quantity of neutrophils. *Oribacterium* genus belongs to the *Lachnospiraceae* family, and the abundance of this genus increased in piglets after weaning [49]. Members of the genus *Oribacterium* produce short-chain fatty acids such as acetate [50], which directly influences immune system regulation [51], and can contribute to the health of the pig.

Among the most connected traits in our network we also found acute-phase protein Hp, which based on the estimated microbiability appeared preferentially determined by microbiota effects. The main function of Hp is to facilitate hemoglobin (Hb) clearance. After the formation of stable Hp-Hb complexes, the macrophage receptor CD163 recognize them and the entire complex is removed from circulation by receptor-mediated endocytosis [52]. Therefore, Hp favors the reduction of free iron concentrations in the circulation and tissues [53]. Several bacterial pathogens such as *Staphylococcus*, *Mycobacteria*, *Salmonella*, *Corynebacterium*, *Haemophilus*, among others, require iron for growth, thus elaborating different acquisition strategies to uptake heme from the host, particularly from Hb [54–56]. The host immune system has developed antimicrobial mechanisms, most related to innate pathways, to deplete iron availability for pathogens [54]. Remarkably, our results indicate a relevant effect of the microbiota composition on Hp levels which could also modulate the concentration of circulating free iron. We could hypothesize that the symbiotic microbiota could also modulate the iron levels in these animals through innate immunity mechanisms to prevent the development of different opportunistic pathogens. Our results confirmed a positive association between serum concentration of Hp and the relative abundance of *Megasphaera*, a member of the phylum Firmicutes. According to this result, an increase in the abundance of *Megasphaera* has been described in colon content and faeces of pigs fed with iron-deficient diet [57]. Interestingly, this genus was reported as a potential biomarker for immune-mediate mechanism of protection from diarrhea [58] and positive correlated with luminal IgA concentration in pigs [49].

Finally, it is worth highlighting the negative association between the relative abundance of *Desulfovibrio* and LEU and MON counts. *Desulfovibrio* is a sulfate-reducing bacteria (SRB), which can promote the metabolism of sugars [59] and plays also a key role in intestinal hydrogen and sulfur metabolism [60]. In pigs, *Desulfovibrio* plays a relevant role during pig gut colonization [49] and was among the dominant genus in healthy pigs compared with diarrhea-affected piglets [61]. In fact, in weaned piglets, a negative correlation between *Desulfovibrio* and several inflammatory markers such as IL-1β, IL-2 and IL-6, have been observed [62], which would be in agreement with the negative correlation observed between *Desulfovibrio* and LEU and MON counts in our piglets.

Despite of an inventory of potential gut health biomarkers exists for pigs [63, 64], our results propose new microbial candidates, and emphasize a polymicrobial nature of the immunocompetence in pigs. Furthermore, in agreement with previous reports [65], our results suggest that some immunity traits are influenced by specific microorganisms while others are determined by interactions between members of the gut microbiome. We also reveal the joint contribution of the host genome and the gut microbial ecosystem to the phenotypic variance of immunity parameters and advice that ignoring microbiota effects could generate an overestimation of genetic parameters. A better understanding of the host and microbial contribution to immunocompetence will allow to develop holistic breeding strategies to modulate these traits, as well as to improve animal health, robustness and welfare.

## Conclusions

Estimates of heritability and microbiability exposed the joint contribution of both the host genome and the gut microbial ecosystem to the phenotypic variance of immunity parameters, and revealed that ignoring microbiota effects on phenotypes could generate an upward bias in the estimation of genetic parameters. Results from the MWAS suggested a polymicrobial nature of the immunocompetence in pigs and highlighted associations between the compositions of pig gut microbiota and 15 of the analyzed immunity traits. Overall, our findings establish several links between the gut microbiota and the immune system in pigs, underscoring the importance of considering both sources of information, host-genome and microbial level, for the genetic evaluation and the modulation of immunocompetence in pigs.

## Acknowledgements

The authors warmly thank all technical staff from *Selección Batallé* S.A, for providing the animal material and their collaboration during the sampling.

