## Supplementary table 1 for "Leveraging host-genetics and gut microbiota to determine immunocompetence in pigs"

| Trait | Full name | h2 | sd | P(h2>0.1) | m2 | sd | P(m2>0.1) | ILh2G | SLh2G | ILh2M | SLh2M |
|---|---|---|---|---|---|---|---|---|---|---|---|
| CRP | C-reactive protein | 0.142 | 0.046 | 0.900 | 0.180 | 0.059 | 0.942 | 0.073 | 0.254 | 0.068 | 0.291 |
| EOS | eosinophils | 0.186 | 0.061 | 0.962 | 0.207 | 0.063 | 0.986 | 0.082 | 0.316 | 0.094 | 0.331 |
| γδ T-cells | gamma-delta T cells | 0.147 | 0.067 | 0.819 | 0.152 | 0.053 | 0.851 | 0.063 | 0.201 | 0.055 | 0.258 |
| GRANU_PHAGO_FITC | mean fluorescence in FITC among granulocytes | 0.255 | 0.061 | 1.000 | 0.234 | 0.056 | 1.000 | 0.135 | 0.371 | 0.129 | 0.341 |
| GRANU_PHAGO_% | percentage of phagocytic granulocytes | 0.199 | 0.052 | 0.995 | 0.180 | 0.062 | 0.933 | 0.100 | 0.300 | 0.068 | 0.300 |
| HP | Haptoglobin | 0.138 | 0.040 | 0.853 | 0.276 | 0.082 | 0.996 | 0.066 | 0.210 | 0.123 | 0.435 |
| IgA | immunoglobulins IgA | 0.220 | 0.081 | 0.996 | 0.217 | 0.073 | 0.980 | 0.101 | 0.373 | 0.075 | 0.355 |
| IgAsal | IgA saliva | 0.202 | 0.051 | 0.997 | 0.210 | 0.073 | 0.975 | 0.116 | 0.309 | 0.081 | 0.351 |
| IgG | immunoglobulins IgG | 0.316 | 0.092 | 1.000 | 0.176 | 0.059 | 0.939 | 0.147 | 0.486 | 0.074 | 0.293 |
| IgM | immunoglobulins IgM | 0.359 | 0.087 | 1.000 | 0.158 | 0.046 | 0.909 | 0.193 | 0.535 | 0.069 | 0.246 |
| LEU | total number of leukocytes | 0.201 | 0.057 | 0.997 | 0.177 | 0.054 | 0.948 | 0.104 | 0.301 | 0.076 | 0.279 |
| LYM | lymphocytes | 0.194 | 0.055 | 0.972 | 0.214 | 0.063 | 0.985 | 0.091 | 0.294 | 0.094 | 0.331 |
| LYM_PHAGO_FITC | mean fluorescence in FITC among lymphocites | 0.242 | 0.054 | 1.000 | 0.260 | 0.053 | 1.000 | 0.145 | 0.332 | 0.155 | 0.362 |
| LYM_PHAGO_% | pertcentage of phagocytic lymphocytes | 0.207 | 0.051 | 0.991 | 0.238 | 0.066 | 0.998 | 0.111 | 0.298 | 0.109 | 0.365 |
| MON_PHAGO_FITC | mean fluorescence in FITC among monocytes | 0.200 | 0.050 | 0.998 | 0.196 | 0.052 | 0.994 | 0.109 | 0.292 | 0.100 | 0.294 |
| MON_PHAGO_% | pertcentage of phagocytic monocytes | 0.189 | 0.060 | 0.978 | 0.219 | 0.065 | 0.993 | 0.087 | 0.298 | 0.101 | 0.343 |
| MONOCITOS_MM | monocytes | 0.153 | 0.036 | 0.972 | 0.173 | 0.054 | 0.944 | 0.091 | 0.223 | 0.077 | 0.277 |
| NO | Nitric Oxide | 0.211 | 0.066 | 0.988 | 0.231 | 0.078 | 0.983 | 0.092 | 0.349 | 0.086 | 0.377 |
| PHAGO_FITC | mean fluorescence in FITC among all phagocytic cells | 0.256 | 0.064 | 1.000 | 0.220 | 0.055 | 0.999 | 0.134 | 0.381 | 0.113 | 0.322 |
| PHAGO_% | percentage of total phagocytic cells | 0.157 | 0.047 | 0.946 | 0.190 | 0.068 | 0.938 | 0.081 | 0.248 | 0.069 | 0.319 |
| NEU | neutrophils | 0.239 | 0.061 | 1.000 | 0.155 | 0.050 | 0.884 | 0.122 | 0.352 | 0.056 | 0.252 |
