## Supplementary figure S1 for "Leveraging host-genetics and gut microbiota to determine immunocompetence in pigs"

**Supplementary figure S1**. Iris-plot representing the 20 most abundant genera. Each bar represents a sample, and bar colors represented the genera relative abundance.

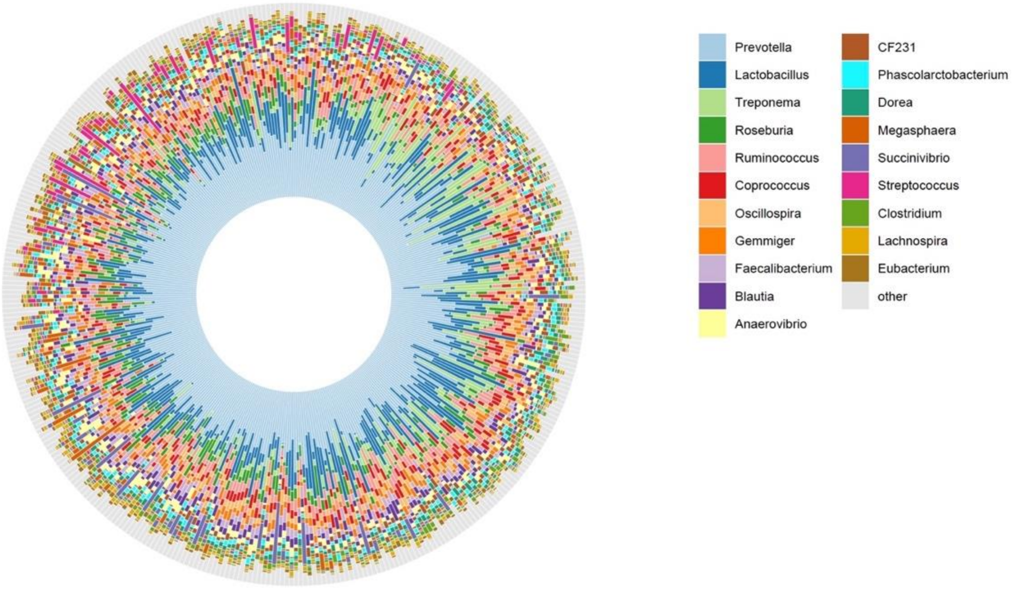
